# Recruitment of Corticotropin-releasing Hormone CRH Neurons in response to Categorically Distinct Stress Challenges in the Mouse Brain

**DOI:** 10.1101/2022.01.20.477065

**Authors:** Krisztina Horváth, Balázs Juhász, Dániel Kuti, Szilamér Ferenczi, Bibiána Török, Dóra Zelena, Krisztina J. Kovács

**Author notes:** Corresponding author: Krisztina J. Kovács, PhD, Laboratory of Molecular Neuroendocrinology, Institute of Experimental Medicine, Szigony u 43. H-1083 Budapest, Hungary.

## Abstract

Corticotropin-releasing hormone (CRH) neurons in the paraventricular hypothalamic nucleus (Pa) are in the position to integrate stress-related information and initiate adaptive neuroendocrine-, autonomic-, metabolic- and behavioral responses. In addition to hypophyseotropic cells, CRH is widely expressed in the CNS, however their involvement in organization of the stress response is not fully understood. In these experiments, we took the advantage of recently available Crh-IRES-Cre; Ai9 mouse line to study the recruitment of hypothalamic and extrahypothalamic CRH neurons in categorically distinct, acute stress reactions. Reporter mice were exposed to restraint, ether, high salt and lipopolisacharide stress. The induced activation of CRH neurons was detected by colocalization of immediate early gene c-Fos in Td tomato expressing cells. We found differential activation of CRH neurons in central amygdaloid nucleus (Ce), bed nucleus stria terminalis lateral division, ventral posterior (BSTLVP), medial preoptic nucleus (MPO), ventromedial hypothalamic nucleus (VMH), premammillary nucleus (PM) and prepositus hypoglossal nucleus (Pr) in response to physiological (ether, high salt and LPS) and psychological (restraint) stressors. CRH positive cells in Pa became activated, however in the Barrington’s nucleus and locus coeruleus no actiovation could be observed due to most tested stressors. Furthermore no CRH neuron activation occured in dorsal (BSTLD) and posterior (BSTLP) region of lateral division of bed nucleus stria terminalis after restraint stress. In the inferior olive only ether exposition resulted in CRH neuron activation. These results confirm activation of CRH neurons in the Pa, reveals new subset of stress-related CRH cells through the mouse brain and disprove the recruitment of CRH cells in the SOL and in the Barrington’s nucleus to acute psychological stress in mice.

## 2. Introduction

Corticotropin-releasing hormone is a 41 amino acid neuropeptide, which plays an essential role in the neuroendocrine stress regulation [1.]. As the key player of this response, it is generally known that the paraventricular hypothalamic nucleus (Pa) integrates external and visceral stress-related informations, and via CRH synthesis and additional factor (e.g., arginine vasopressin (AVP)) release into the hypophyseal portal circulation, triggers and modulates ACTH secretion from the pituitary corticotropes, which then stimulates corticosteroid discharge from the adrenal cortex [2.]. In addition to the Pa, CRH expressing neurons are widely found in the CNS, where they might serve neurotransmitter and neuromodulatory role. While the structure and function of Pa CRH neurons are well established, much less is known about the role and organization of CRH cells outside of the hypothalamus. Detecting and visualizing CRH-positive neurons in the brain is challenging due to its low basal level in the cell bodies and the absence of highly sensitive CRH antibodies. Early studies used in situ hybridization to provide information about the localization of CRH mRNA expression. The drawback of the method is that regions which are rich in nuclei often have high backgrounds and are easily mistaken as positive signals. So justification of the labeling actuality is critical [3.]. CRH positive neurons can also be detected by immunohistochemistry with invasive tissue preparation [4.], although with this method the amount of revealable cells is limited [5.]. Another well-known way to identify CRH neurons is colchicine treatment. Adminidtration of the medication results in the accumulation of the peptid in soma of neurons by blocking the axonal transport. Dispite elevating the immunoreactivity of CRH cells, physiological relevancy of the detected nuerons are doubtful, as colchicine treatment can evoke altertion of neuronal regulation itself [6.]. A recently developed technique, crossing CRH-Ires-CRE mice with Cre-reporter mice, allowes direct visualization of CRH neurons. Besides no secondary detection method is needed using these animals, genetic modification has no act on the physical state of the mice. Furthetmore, it is notable, that while with immunohistochemical staining a current protein content in a certain timepoint above a detection threshold can be identified, with genetic modification tracking of CRH expression over the whole lifespan becomes possible [7.]. Therefor with this pattern, detailed information can be mined upon the exact location of CRH neurons in distinct regions of the mouse brain.

In our experiment we prepared a whole brain CRH distribution analysis on Crh-IRES-Cre;Ai9 mice. In order to determine the potential discrepancy in the activation of each found population due to physiological and psychological acute stressors, c-Fos analysis was performed. Immediate early gene (IEG) c-Fos, is rapidly and transiently induced by various neuronal inputs, which results in activation in the target cells [8.]. Immunocytochemical detection of the IEG protein product c-Fos has long been used as a functional anatomical marker to visualize activated neurons by different stress challenges [9.]. Overall, with the available detection facilities, the probable differences in stress response paradigm of variant stressors can be examined.

## 3. Materials and methods

### 3.1. Animals

Crh-IRES-Cre (B6(Cg)-Crh^tm1(cre)Zjh^/J; stock number: 012704) and Ai9 mice (B6;Cg-Gt(ROSA)26Sor^tm9(CAG-tdTomato)Hze^/J; stock number: 007905) were purchased from Jackson Laboratory. The used Crh-IRES-Cre;Ai9 animals −in which Cre mediated recombination [10.] resulted in tdTomato fluorophore expression in Crh neurons −were derived from second crosses of homozygous Crh-IRES-Cre/Ai9 genotypes. Mice were bred at the specific pathogen free (SPF) level and were maintained at the minimal disease (MD) level of the Transgenic Facility of the Institute of Experimental Medicine, Hungary. In our experiments adult mice (10-12 weeks of age) were used. Animals had ad libitum access to standard laboratory animal pellet and water. They were kept under the circumstances of 21±1 °C, 400 lx light intensity, 65% humidity and 12-12 hour light-dark phase. The procedures which we used in the experiment were in accordance with the guidelines set by the European Communities Council Directive (86/609 EEC) and approved by the Institutional Animal Care and Use Committee of the Institute of Experimental Medicine.

### 3.2. Acute stress

Crh-IRES-Cre;Ai9 mice were caged individually one day prior the stress treatments. Animals were exposed to restraint (n=6), ether (n=3), lipopolysaccharide (LPS) (n=3) or hypertonic saline (n=2) stress. Tests were carried out in a separate experimental room in the early light phase of the day. Light intensity, humidity and temperature were analogue to that in the maintenance room. After the exposition animals were placed back to their homecage until sacrifice.

#### 3.2.1. Restraint stress

Restraint stress was performed using transparent 50 ml Falcon tubes, in which holes were cut at the end and alongside the plastic to prevent overheating of the animals. Mice were captured in the tubes for 30 minutes. To achieve comparable degree of restraint, packing with paper towels at the rear was used. This procedure minimized the space around the animal, prevented them from turning and provided stressful stimulus, without being harmful.

#### 3.2.2. Ether stress

To accomplish the ether stress, mice were placed into a lockable glass pot in which ether soaked cotton was placed along with a paper towel to prevent the fur of the animals from soaking. After dormition mice were removed from the bottle and the etheric anaesthesia was maintained up to 5 minutes with a nosecone filled with additional ether soaked cotton.

#### 3.2.3. Lipopolysaccharide stress

Lipopolysaccharide (serotype: 0111:B4, Sigma L4391) was administered intraperitoneally at dose of 1 mg/ml (0.1 ml/100g). The dose was selected to induce substantial systemic inflammation, without compromising survival (LD50 value for LPS in mice after intraperitoneal administration is 10-20 mg/kg).

#### 3.2.4. Hypertonic saline stress

Mice were injected intraperitoneally with hypertonic saline solution (1.5M, 1.8 ml/100 g).

#### 3.2.5. Controls

For the the restraint and ether stressed animals, undisturbed mice were considered as control group (n=6). In case of lipopolysaccharide and hypertonic saline stressed animals, controls were injected intraperitoneally with physiological saline solution (0.1 ml/100g) (n=2).

### 3.3. Tissue processing

90 minutes after the beggining of the stress expoure, under terminal anesthesia (Nembutal, Ceva-Phylaxia, Budapest, Hungary) animals were transcardially perfused with 0.9% saline followed by 70 ml ice-cold fixative (4% paraformaldehyde in 0.4 M phosphate buffer, pH 7.2). Brains were removed and postpost-fixed in the same fixative complemented by 10% sucrose for 3 hours and cryoprotected overnight in 10% sucrose in potassium phosphate buffered saline, KPBS. Four series of coronal sections (25 μm) were cut on freezing microtome. Sections were stored at −20 °C in antifreeze solution (30% ethylene glycol and 20% glycerol in 0.1M PBS).

### 3.4. c-Fos immunocytochemistry

Free-floating brain sections were incubated in 2% normal goat serum (Vector Laboratories, Burlingame, CA) in PBS/0.3% Triton X100 at room temperature for 1 hour; rabbit anti-c-Fos IgG (sc-52 Santa Cruz Biotechnology, Santa Cruz, CA, 1:20000) at 4 °C overnight; Alexa Fluor 488 donkey anti-rabbit IgG (ThermoFisher, 1:500 dilution) at room temperature for 2 hours in dark. Sections were mounted onto slides and coverslipped with DAPI Fluoromount-G (Southern Biotech).

### 3.5. Imaging, quantification and data analysis

Digital images of the brain sections were captured at 20x magnification with 3D HISTECH Pannoramic MIDI II. slide scanner. Regions of interest were outlined on the basis of Allen Brain Atlas and then were analyzed with NIS Elements Imaging Software 5.21.01. Number of DAPI, tdTomato and c-Fos positive cells and total area were captured of each region. Correlation amongst number of DAPI positive cells and area size of the region of interest was examined. CRH neuron ratio was counted as number of the tdTomato positive cells divided by the total cell amount of the specific region*100. Activated neuron ratio was counted as number of the c-Fos positive cells divided by the total cell amount of the specific region*100. Treatment induced differences was calculated as treated animal [number of CRH or c-Fos positive cells divided by the total cell amount of each region*100] - control animal [number of CRH or c-Fos positive cells divided by the total cell amount of each region*100]. Density of neurons was defined as the signaling cell number divided by area of the specific region. Outcome of the restraint represented the effect of psychological, and outcome of high salt, ether and LPS administration represented the effect of physiological stress.

### 3.6. Corticosterone measurement

After decapitation, trunk blood was collected into plastic tubes filled with 10 μl of 20% K-EDTA. Samples were centrifuged and plasma were stored at −20 °C. Corticosterone was determined using a direct radioimmunassay (RIA) as described [11.].

### 3.7. Statistical analysis

Data were analyzed using GraphPad Prism software (version 7; San Diego, CA, USA). Correlation analysis (comparison of DAPI positive cell number to the area size), one-way ANOVA with Tukey’s multiple comparisons test (CRH neuron distribution; neuronal activation and colocalization analysis) and unpaired t-test (determination of discrepancy between psychological and physiological stressed animals) were performed. We considered p-value<0.05 as statistically significant. Data are expressed as group mean ± SEM (standard error of the mean) for each treatment group.

## 4. Results

### 4.1. Correlation analysis

The total cell amount and area of the region of interest significantly correlated (p<0.0001).

### 4.2. CRH neuron distribution

By the examination of 95 brain region, no significant difference could be detected in average CRH ratio among the experimental animals. Further monitoring tdTomato positive neuron distribution showed that the greatest average ratio of CRH cells occures in the brainstem, striatum and limbic area with 6.60%, 6.33% and 6.28%. Also greater cell rate could be monitored in thalamus, midbrain, hypothalamus and cortical plate regions. Average distribution in these areas were 3.35%, 3.27%, 3.18% and 3.03%. As further disassembling hypothalamus region, periventricular zone proved to be the most possessed by tdTomato positive neurons with an average of 8.52%, while periventricular region, hypothalamic lateral zone and hypothalamic medial zone harboured less CRH neurons: 2.82%; 2.12% and 2.10%. In case of the cortical plate, larger tdTomato ratio could be monitored in the olfactory area and the isocortex with an average of 3.75% and 3.17%. Examination of the hippocampal formation showed an average of 0.67% CRH cell ratio. In point of the 95 counted area, more regions are worthful to be mentioned regarding to their high average CRH neuron density. In the brainstem, IO harbored 25.76%, Bar harbored 16.53%, X harbored 13.24%, LRt harbored 11.64%, Cu harbored 8.28%, Rpa harbored 7.50%, DMPn harbored 6.77%, Ecu harbored 5.86% and MVePC harbored 5.10%. In striatum two regions were outstanding. HDB with 16.41% and Ce with 11.38%. In the limbic area almost all examined area showed higher than an average of 5% CRH positive cell count. BAC showed 9.61%, BSTLVP showed 7.25% and BSTLP showed 6.26%. In the thalamus there were two prominent region: PVP with 7.12% and PIL with 5.26%. Overall the second most tdTomato positive cell harboring area could be found in the periventricular zone of the hypothalamus. Unlike the other regions in this field, Pa showed high, 23.51% ratio. In the midbrain IPAC and PrC showed the highest, 6.94% and 5.78%. In the olfactory area EPIO with 13.92%, Pir with 5.69%, and in the isocortex region PrL with 6.28% ensured the high average tdTomato positive cell density in the cortical brain region.

### 4.3. Neuronal activation

#### 4.3.1. Restraint

Examining the average discrepancy from the control group, restraint induced more neuronal activation in the hypthalamus with 5.91%. This value in the thalamus was 5.45%, in the cortical plate 4.92%, in the limbic area 3.94% and in the striatum 3.81%. Compared to the hypothalamic area, midbrain with 2.84% (p=0.0021) and braintstem with 2.05% (p<0.0001) showed significantly lower c-Fos expression. Number of signaling cells in the midbrain (p=0.0120) and brainstem (p=0.0005) remained on a significantly lower level compared to the thalamus. In case of the cortical plate, mostly the olfactory area and isocortex showed considerable 7.05% and 4.67% elevation. In the hippocampal formation 1.31% increase colud be monitored. In the hypothalamic region, mainly periventricular zone and hypothalamic medial zone with 11.7% and 8.21% enhanced the average ratio. While in the hypothalamic lateral zone 4.01%, in the periventricular region 1.73% growth could be observed. In detail, significant neuronal activation discrepancy could be monitored due to restraint stress in GrO, VTT, DP, DEN, Pir, AI, O, LVOV, S1, Cl, GI, PrL, IL, Cg, Aco, LOT, LS, Me, BSTLD, BSTLP, BSTLVP, Rh, PVA, PMTN, IAD, PIL, G, VMPA, MPO, SCh, LH, AHP, PeF, MCLHV, Pa, VMH, DM, PHD, PHV, PH, Su, PrC, PAG, MnR, CG, Bar and Sol.

#### 4.3.2. Ether

Compared to the control group, ether stressed animals displayed the biggest alteration in average neuronal activation in the cortical plate with 9.51% and in the thalamic area with 9.00%. In the hypothalamus 5.86%, in the striatum 4.70%, in the midbrain 3.50%, in the limbic area 3.46% and in the brainstem 1.82% elevation could be seen. The average activation in the thalamus was significantly higher compared to the striatum (p=0.0007), limbic area (p<0.0001), brainstem (p<0.0001), midbrain (p<00001) and hypothalamus (p=0.0118). Deviation was also significant between cortical plate and striatum (p=0.0002), limbic area (p<0.0001) brainstem (p<0.0001), midbrain (p<0.0001) and hypothalamus (p=0.0034). Furthermore the difference between hypothalamus and brainstem was also significant (p=0.0013). Overall, in the cortical plate, the olfactory area with 13.09% and the isocortex with 9.26% ensured the higher average areal activation. Hippocampal formation showed only 2.48% increase. In case of the hypothalamic lateral zone, 10.83% rising could be observed. This value in the hypothalamic medial zone was 6.8%, in the hypothalamic lateral zone 4.75% and in the periventricular region 2.72%. Analysation of each specific region showed significant shift of c-Fos activation in GrO, VTT, DP, DEN, Pir, M2, Al, M1, O, DTr, LVOV, S1, Cl, GI, PrL, IL, Cg, Aco, LOT, RS, HC, DG, CA3, Acb, Ce, Me, BSTLD, BSTLP, BAC, Rh, PVA, CM, PMTN, IAD, PIL, G, VMPA, MPO, LH, AHP, STh, MCLHV, ZI, Pa, Arc, VMH, PHD, PHV, PH, Su, IPAC, PrC, EW, IPR, CG, Cu, AP and Sol compared to the control group.

#### 4.3.3. High salt

High osmolarity stress induced the elevation of average neuronal activation compared to the control group in the limbic area with 6.89%, in the hypothalamus with 5.63%, in the thalamus with 5.41%, in the cortical plate with 4.83%, in the striatum with 3.85%, in the midbrain with 3.00% and in the brainstem with 2.82%. In detail, olfactory area showed 7.19%, isocortex 4.71% and hippocampal formation 0.51% increase. In the hypothalamus, the most altered activational regio was the periventricular zone with 16.66% increase. Hypothalamic medial zone showed only 4.47%, hypothalamic lateral zone 3.87% and periventricular region 2.9% growth. Regarding the 95 smaller areas, as result of high osmolarity stress, significant alteration could be detected in the number of activated neurons compared to the control group in the region of AI, M1, Ce, Me, BSTLVP, VMPA, PeF, PSTh, Pa, IF, X and Sol.

#### 4.3.4. LPS

Compared to the control group, LPS treated animals showed an average of 6.83% neuronal activation growth in limbic area. Additional average elevation could be monitored in the hypothalamus with 4.45%, in the thalamus and midbrain with 4.19%, in the brainstem with 3.31%, in the striatum with 2.69% and in the cortical plate with 2,67%. Further diassembly of cortical plate showed that isocortex played the largest role in the above mentioned data with 15.51%. In case of olfactory area and hippocampal formation, the rate of growth was only 4.30% and 0.93%. In the hypothalamic area periventricular zone showed 15.22% increase. Hypothalamic lateral zone fell behind with 2.9%, hypothalamic medial zone with 2.67% and periventricular region with 2.6% elevation. Compared to the control group, LPS caused the change of c-Fos positive cell ratio significantly in AI, M1, Ce, Me, BSTLP, BSTLVP, VMPA, PeF, PSTh, Pa, IF, X and Sol.

### 4.4. Colocalization

#### 4.4.1. Restraint

Acute stress resulted in the alteration of average C-Fos and tdTomato positive cell colocalization in several brain regions. At the greatest rate, restraint effected the hypothalamic area, in which an average of 0.89% increase colud be observed. In the limbic area this ratio dropped to 0.44%, in the cortical plate to 0.26%, in the brainstem to 0.25%, in the thalamus to 0.19%, in the midbrain to 0.17% and in the striatum to 0.14%. Colocalization in the hypothalamus was significantly higher compared to that in the cortical plate (p=0.0170), in the brainstem (p= 0.0160), in the thalamus (p=0.0065), in the midbrain (p=0.0045) and or in the striatum (p=0.0031). Olfactory area, isocortex and hippocampal formation shared an average of 0.38%, 0.24% and 0.04% increase. From the point of hypothalamus, the greatest elevation could be observed in the periventricular zone with 5.30%. In the hypothalamic lateral zone 0.24%, in the hypothalamic medial zone 0.14% and in the periventricular region 0.13% growth could be monitored. Overall, restraint induced significant colocalization alteration in AO, DP, Pir, AI, O, LVOV, Cg, HDB, BSTLVP, PMTN, MPO, AHP and Pa compared to the control group.

#### 4.4.2. Ether

As an aftermath of ether exposition, average colocalization increase could be measured in the hypothalamic area by 0.71%, in the cortical plate by 0.56%, in the limbic area by 0.52%, in the thalamus by 0.34%, in the striatum by 0.28% and in the brainstem by 0.22%. In contrast, midbrain showed an average of 0.03% decrease. The difference in the change rate was significant between hypothalamus and striatum (p=0.0440), brainstem (p= 0.0200) and midbrain (p=0.0005). The same footage could be seen in case of midbrain and cortical plate (p=0.0044) and limbic area (p=0.0074). Further division showed that olfactory area went through a 0.71%, isocortex a 0.59% and hippocampal region a 0.06% growth. In the hypothalamus, periventricular zone showed 4.41%, hypothalamic lateral zone showed 0.17% and hypothalamic medial zone showed 0.11% and periventricular region showed 0.08% rate growth. Compared to the control group, further colocalization discrepancy could be monitored in EPIO, DP, Pir, M2, AI, O, LVOV, S1, Gl, IL, Cg, LOT, RS, DG, Ce, BSTLD, CM, PMTN, MPO, Pa, PM, Cu and IO.

#### 4.4.3. High salt

High osmolarity stress induced alteration in the rate of average colocalization in several areas compared to the control group. Hypothalamus showed 0.88%, brainstem showed 0.6%, thalamus showed 0.49%, cortical plate showed 0.29%, midbrain showed 0.24%, striatum showed 0.23% and limbic area showed 0.21% increase. Hypothalamic growth proved to be significant compared to that in the cortical plate (p=0.0372), midbrain (p=0.0249), striatum (p=0.0229) and limbic area (p=0.0187). In the elevation of colocalization rate in the cortical plate, isocortex and olfactory area played nearly equal role with 0.34% and 0.33%. Hippocampal formation showed 0.01% rising. Colocalization rate of periventricular zone emerged from that in the other hypothalamic regions with 5.35%. Hypothalamic lateral zone showed 0.26%, hypothalamic medial zone showed 0.16% and periventricular region showed 0.04% elevation. Further significant deviation could be noted in Cl, Ce, BSTLD, CM, PVP, VMPA, Pa, PHV, PrC and RtTg compared to the control group.

#### 4.4.4. LPS

Changing of the average colocalization rate could be observed in distinct regions of the mouse brain in case of LPS stressed animals compared to the control group. The graetest rate growth was 0.76% which could be monitored in the hypothalamic region. This was fallowed by the brainstem with 0.32%, limbic area with 0.2%, thalamus with 0.17%, cortical plate with 0.14%, striatum with 0.13% and midbrain with 0.01%. Significant difference could be measured between the increase rate of hypothalamus and cortical plate (p=0.3950), striatum (p=0:0357) and midbrain (p=0.0105). Examining the cortical plate, isocortex showed 0.15%, hippocampal formation 0.14% and olfactory area 0.12% increase. In case of the hypothalamus, periventricular zone showed the largest 5.18% elevation. Hypothalamic lateral zone and hypothalamic medial zone showed only 0.16% and 0.02% increase. Periventricular region endured an average of 0.05% decrease in colocalization rate. Among Ce, BSTLP and Pa further significant alterations could be observed compared to the control group.

### 4.5. Stress effect differences

Comparing the average neuronal activation among treated groups, several significant discrepancies could be monitored. The shift of c-Fos positive cell count from the control group was significantly higher in the restraint stressed animals compared to the LPS treated mice in AL and PeF. Furthermore ether exposition elevated neuronal activation more in DP, Pir, M2, DTr, PrL, IL, Ce, PM, IPAC and Cu compared to the restraint group. The same was noted towards high osmolarity stress in PrL, IL and Rh. Also bigger grwoth was detectable in DP, Pir, M2, Al, PrL, IL, Rh, IAD, MPO, PM and IPR compared to LPS treatment. In case of high salt stress, the increase of c-Fos positive cells ratio proved to be significantly higher in Ce, BSTLVP, SO and Sol compared to the restraint stressed mice. They also expressed elevated ratio compared to the ether exposed animals in BSTLVP, SO and Sol. Similar discrepancy was observable from the LPS group in the Al region. As a consequence of LPS injection, more neuron activated in Ce, BSTLP, BSTLVP, SO and Sol compared to the restraint group. This effect was also observable contrast to the ether exposed animals in BSTLP, BSTLVP and Sol.

In point of colocalization ratio growth compared to the control group, restraint stressed mice showed significantly greater elevation in MPO in contrast to the ether exposed mice. Also higher rising was detectable in the restraint group compared to the high osmolarity stressed animals in MPO and Pr. The same was observable compared to the LPS treated animals in MPO, VMH and Pr. Ether exposition resulted in significantly higher colocalization rate growth in M2, Ce, PM and Cu regions in contrast to the restraint stress. It also provoked significantly higher colocalization elevation in M2, PM and Cu than high salt treatment. In M2, Al, PM and Cu ether stressed animals showed greater colocalization ratio rising compared to the LPS group. High osmolarity stress lifted the colocalization significantly more in the LRt compared to the ether exposition. LPS evoked significantly more colocalization in the C than ether exposition.

Examination showed significant differences in the alteration of neuronal activation and colocalization due to psychological and physiological stress. As an aftermath of restraint, significant elevation of neuronal activation colud be detected in MnR, PeF, PrC, DM, LH, Su and PH compared to the animals of the ether, high salt or LPS treated group. In contrast, physiological stress resulted in significantly higher c-Fos positive cell ratio increase in Ce, Sol, SCh, BSTLP, IPAC and SO than psychological stress. In point of colocalization, psychological stress caused significant elevation in BSTLVP, MPO, VMH and PM, while physiological stress in Ce and Pr compared to each other.

## 5. Conclusion

It is well understood that acute stress besides activating the sympathetic nervous system, evokes neuroendocrine stress response by disturbing homeostasis [12.]. In our experiment the activation of hypothalamic-pituitary-adrenal (HPA) axis could also be detected, as each stressors resulted in robust neuronal activation in the Pa, from which more than 60% were CRH cells. As a further general confirmation of the forming stress cascade is elevated corticosterone level, which was also confirmed to rise in case of several acute homeostatic challenges [13., 14., 15.]. While the main participators and process of stress paradigm is established, it is not clearly known how additional CRH neuron populations throughout the mouse brain affect this mechanism. Furthermore as different types of stressors may not necessarily evoke parallel changes in biochemical and physiological processes [16., 17.], the bias in the exact role of the clusters also still remains unknown. Here we have shown that, the lately popular genetically modified Crh-IRES-Cre;Ai9 mice after being exposed to variant acute stressors, can be successfully used to study the emerged issues. After restraint, ether, high salt or LPS stress, 95 specific brain regions were selected for further analysation according to the ratio of attending activated and tdTomato positive neurons. As validation of CRH cell identification, reliable overlap between transgene expression and CRH mRNA was confirmed by Dr. TZ Baram laboratory [18.].

To monitor the recruitment of CRH cells in response to variant stressors, immediate-early gene activation marker, c-Fos expression analysis was performed. The same anatomic approach has been used in a report by Walker et al. [19.] The timecourse of stress-induced c-Fos activation was comparable to that seen in non-transgenic mice, in principle with maximal c-Fos induction seen at 90 minutes, which is followed by a decline at 3-4 hours post-stress [20., 21., 22.]. However in favour of more specific details, it could be advisable to determine the exact timecourse of c-Fos expression in case of each stressors and brain regions, as due to literature bias can be monitored in homaostatic challenge induced neuronal activation in some cases. For instance in rats biphasic c-Fos induction was found by Ericsson et al. following systemic interleukin-challenge [23.], while Rivest et al. captured delayed elevation of c-Fos mRNA level after LPS injection [24.]. Also delayed signal could be seen in some brain regions [25.]. The result of our measurements showed significant alteration in CRH and additional neuronal activation rate due to the four different stressors compared to the control groups. All together, restraint and ether exposition resulted in greater c-Fos activation, than high salt or LPS treatment. Referring to Meijer [26.], the smaller elevation rate from the control group in the latter cases could be explained by the fact, that intraperitoneal saline injection itself also resultes in a mild stress response. Neuronal activation also showed interesting alteration in some known stress related regions. Aside from the Pa, amygdala has one of the most important role in HPA axis coordination, however the subnuclei of the region serve different functions. While Me has shown to have higher responsiveness to psychogenic stressors, Ce is only crucial to systemic challenges [27.]. The exclusive function of Ce and the prominent role of Me in restraint was also confirmed in our experiment. It is notable however, that while in Ce mainly activation of CRH cells, in Me other cell types could be detected. VMPA and DM stimulation has former been proved to inhibit HPA axis [28.]. In our case VMPA showed significant c-Fos activation elevation in every, DM only in case of restraint stress. From the point of CRH activation, among these areas only VMPA showed elevated signal by high salt stress. In spite of the fact, that on the basis of literature, NTS showes stressor specificity – as neuronal activation tought to only occure after homeostatic challenges, but not after psychogenic ones [29.] –, here all stressors resulted in significant neuronal activation of the region. Only the rate of the elevation was lower in case of restraint stress compared to the other groups. Yet no significant CRH activation elevation could be monitored as a result of any stressors. Other typical examined regions in connection to acute stress are the IL, Prl and the hippocampus. Due to previous research, damage of these regions leads to elevated corticosterone and ACTH responses after restraint stress, but not after ether exposition [30., 31.]. In our case, significant neuronal activation rate growth could be monitored in IL and PrL as result of both restraint and ether stress, while in hippocampus the growth could be detected only after ether stress [32., 33.]. In contrast to this, CRH activation elevation could have been monitored only in IL and in DG part of the hippocampus. PVA also showed significant neuronal, but not CRH cell activation elevation due to restraint and ether exposition, although the area is less known about its effect in acute, but more known to be involved in chronic stress as a prohibitor in habituation of HPA axis responses to stress [34.]. In common, all previously mentioned regions has a connection to the BST which has a shared role in HPA axis regulation and have long been implicated as a relay limbic information to the hypotahalamic neurosecretory neurons [35.]. It is known that divisions of the region has converse function in stress adaptation. While anteriormedial divisions enhances, posterior and anteroventral nuclei inhibit stress axis [36., 37.]. Our experiment also confirmed stressor specificity in neuronal activation of the region. Dorsal and posterior part of lateral division showed significant elevation due to restraint and ether exposition, while in case of ventral posterior region all three used stressors resulted in significant growth except for the ether treatment. Interestingly, despite this, in the latter brain region only restraint showed significantly elevated CRH cell activation.

According to our results, the most tdTomato positive cell dense regions are IO, Pa, Bar, X, LRt, Cu, Rpa, DMPn, Ecu, MvePC, HDB, Ce, BAC, BSTLVP, BSTLP, PVP, PIL, IPAC, PrC, EPIO, Pir and PrL. Similar findings can also be found in the literature [4., 38., 39.]. Examining these regions from the point of CRH cell activation, further functional discrepany can be presumed in the role of the clusters in case of the four different stressors. While the activation rate of Pa CRH cells elevated due to each stressors, Ce – as described before – only showed this paradigm due to physiological stress. CRH neurons in IO, Cu and EPIO only displayed elevated activation after ether, in HDB and BSTLP only after restraint and in PVP and PrC only after high osmolarity stress. In Pir significantly higher colocalization rate growth could be seen in case of restraint and ether treatment. Monitoring the other listed regions showed no significat CRH cell activation elevation due to the applied stressors. This observation leads to the question, whether these cells have any role in stress response, and if not, than what other function might they be responsible for. Some findings have already been published dealing with this question. For instance IO explored to be fundamental in challenge-induced motor responses [40.]. MvePC nuclei is proven to has a strong effect on the HPA axis through vestibulo-paraventricular polysynaptic pathway [41.], withwhich it can modulate or evoke stress response. Due to the literature, Ecu may has a role in the regulation of motor functions [42.]. Barrington’s nucleus - the pontine micturition center [43.] - have previously been implicated in stress-induced urination and its CRH mRNA levels are shown to increase after stress. Despite this, in our experiment no significant colocalization of TdTomato and c-Fos have been found in Bar. Wang et al [44.] described a potential cause that could underlie the function discrepancy of CRH cells: neurons have different dendritic morphologies and projection fibers in distinct brain regions. Some stress response differences although could be derived from other reasons, like specificity of a certain species or aging. Deviation was explored in distribution and density of CRH-immunoreactive neurons between examined mouse and rat [45.]. Altered stress response also can be derived by the changing of CRH producion through the lifecircle [46.].

Further differences could be monitored among the four stressors in regions that mediate mostly the sensory, motor or autonomic functions. For istance, while ether exposition elevated neuronal acitivation and colocalization rate considerably in olfactory area, LPS treated animals fell behind from the other group averages in this regard. High salt administration and LPS injection raised the c-Fos positive neuron rate in the limbic area more than in the other two treatments. In contrast, colocalization rate elevated in the other two groups more, while in case of high salt and LPS treatment remained lower. Neuronal activation in the thalamic region was higher in case of ether administration, while colocalization rate grew most in case of high salt exposition. Neuronal activation due to high salt and LPS also fell behind in periventricular and hypothalamic medial zone. High salt in midbrain and brainstem, while restraint in midbrain elevated the colocalization rate. BSTLVP and Ce nearly showed opposed neuronal activation and colocalisation rate change.

In summary, CRH-Ires-Cre mice crossed with a Cre-dependent reporter strain, represent a useful model in which the neurobiology of the CRH system can be studied. These functional results provide a strong basis for further studies in which the brain CRH recritment should be systematically probed by using opto- and chemogenetic tools to examine the exact role of the differently activated non CRH nuerons in case of distinct stressors. Furthermore it is appropriate to gain information on the function of the other CRH clusters which do not seem to be part of the above mentioned stress responses.

## Abbreviations

Acb: accumbens nu
Al: agranular insular cortex
Aco: anterior cortical amygdaloid nucleus
AHC: anterior hypothalamic area, central
AHP: anterior hypothalamic area, posterior
AO: anterior olfactory nucleus
Arc: arcuate hypothalamic nucleus
AP: area postrema
Bar: Barrington’s nucleus
BAC: bed nucleus anterior commissure
BST: bed nucleus stria terminalis
BSTLD: bed nucleus stria terminalis, lat div, dors
BSTLP: bed nucleus stria terminalis, lat div, post
BSTLVP: bed nucleus stria terminalis, lat div, vent posterior
CA3: CA3 field, hippocampus
Ce: central amygdaloid nucleus
CG: central gray
CM: central medial thalamic nucleus
Cg: cingulate cortex
Cl: claustrum
C: cochlear nucleus
Cu: cuneate nucleus
DpMe: deep mesencephalic nucleus
DG: dentate gyrus
DEN: dorsal endopiriform nu
10: dorsal motor nucleus vagus nerve
DP: dorsal peduncular cortex
DTr: dorsal transition zone
DM: dorsomedial hypothalamic nucleus
DMPn: dorsomedial pontinue nucleus
EW: Edinger-Westphal nucleus
Ecu: external cunate nucleus
EPIO: external plexiform layer, olfactory bulb
G: geniculate nucleus
GI: granular insular cortex
GrO: granule layer, olfactory bulb
HC: hippocampus
HDB: horizontal diagonal band
12: hypoglossal nucleus
IO: inferior olive
IL: infralimbic cortex
IAD: interanterior dorsal thalamic nucleus
IF: interfascicular nucleus
IPR: interpeduncular nucleus, rostral subnucleus
IPAC: interstit nucleus post limb ant com
LVOV: lateral and ventral orbital cortex ventral part
LH: lateral hypothalamic area
LRt: lateral reticular nucleus, ventrolateral medulla
LS: lateral septal nucleus
LC: locus coeruleus
MCLHD: magnocellular nucleus, lateral hypothalamic area dorsal part
MCLHV: magnocellular nucleus, lateral hypothalamic area ventral part
Me: medial amygdaloid nucleus
MPO: medial preoptic nucleus
MVePC: medial vestibular nucleus, parvicellular
MnR: median raphe nucleus
LOT: nucleus lateral olfactory tract
X: nucleus X
O: orbital cortex
PSTh: parasubthalamic nucleus
Pa: paraventricular hypothalamic nucleus
PVA: paraventricular thalamic nucleus, anterior
PVP: paraventricular thalamic nucleus, posterior
PAG: periaqueductal gray
PeF: perifornical nucleus
Pir: piriform cortex
Pn: pontine nuclei
PHD: posterior hypothalamic area, dorsal part
PH: posterior hypothalamic area, posterior
PHV: posterior hypothalamic area, ventral part
PIL: posterior intralaminar thalamic nucleus
PMTH: posteromedial thalamic area
PMTN: posteromedial thalamic nucleus
PrC: precomissural nucleus
PrL: prelimbic cortex
PM: premammillary nucleus
Pr: prepositus hypoglossal nucleus
M1: primary motor cortex
Rpa: raphe pallidus nucleus
RtTg: reticulotegmental nucleus pons
RS: retrosplenial cortex
Rh: rhomboid thalamic nucleus
M2: secondary motor cortex
S2: secondary somatosensory cortex
Sol: solitary tract nucleus
S1: somatosensory 1
STh: subthalamic nucleus
SC: superior colliculus
SCh: suprachiasm nucleus
SO: supraoptic nucleus
Su: suprmammillary nucleus
VTT: ventral tenia tecta
VLG: ventrolateral genic nucleus
VMH: ventromedial hypothalamic nucleus
VMPA: ventromedial preoptic area + OVLT
ZI: zona incerta

## 4.1. Correlation analysis

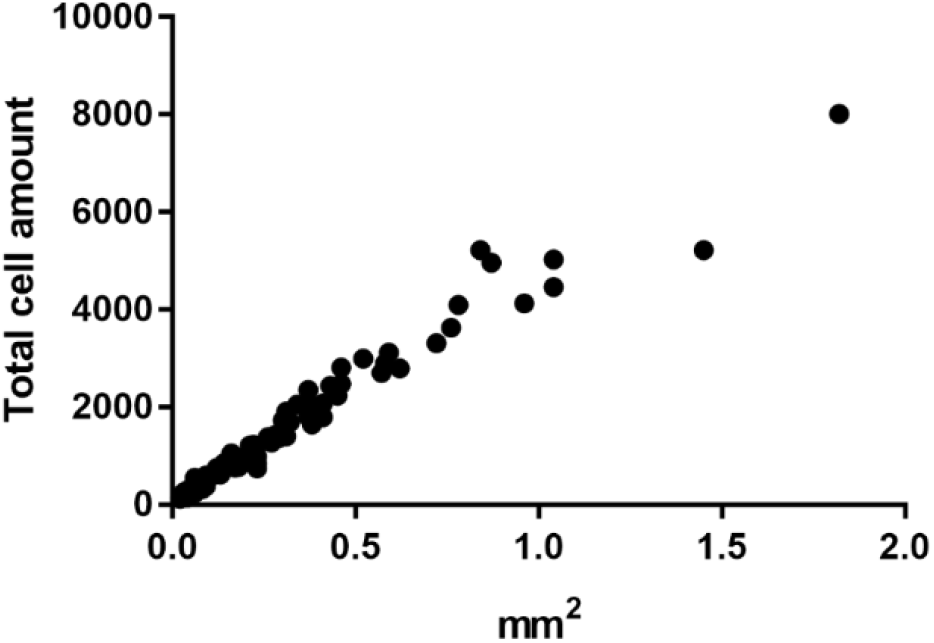
Correlation of average cell amount (number of DAPI positive cells) and averge field area of the 95 examined brain region (n=22).

## 4.2. CRH neuron distribution

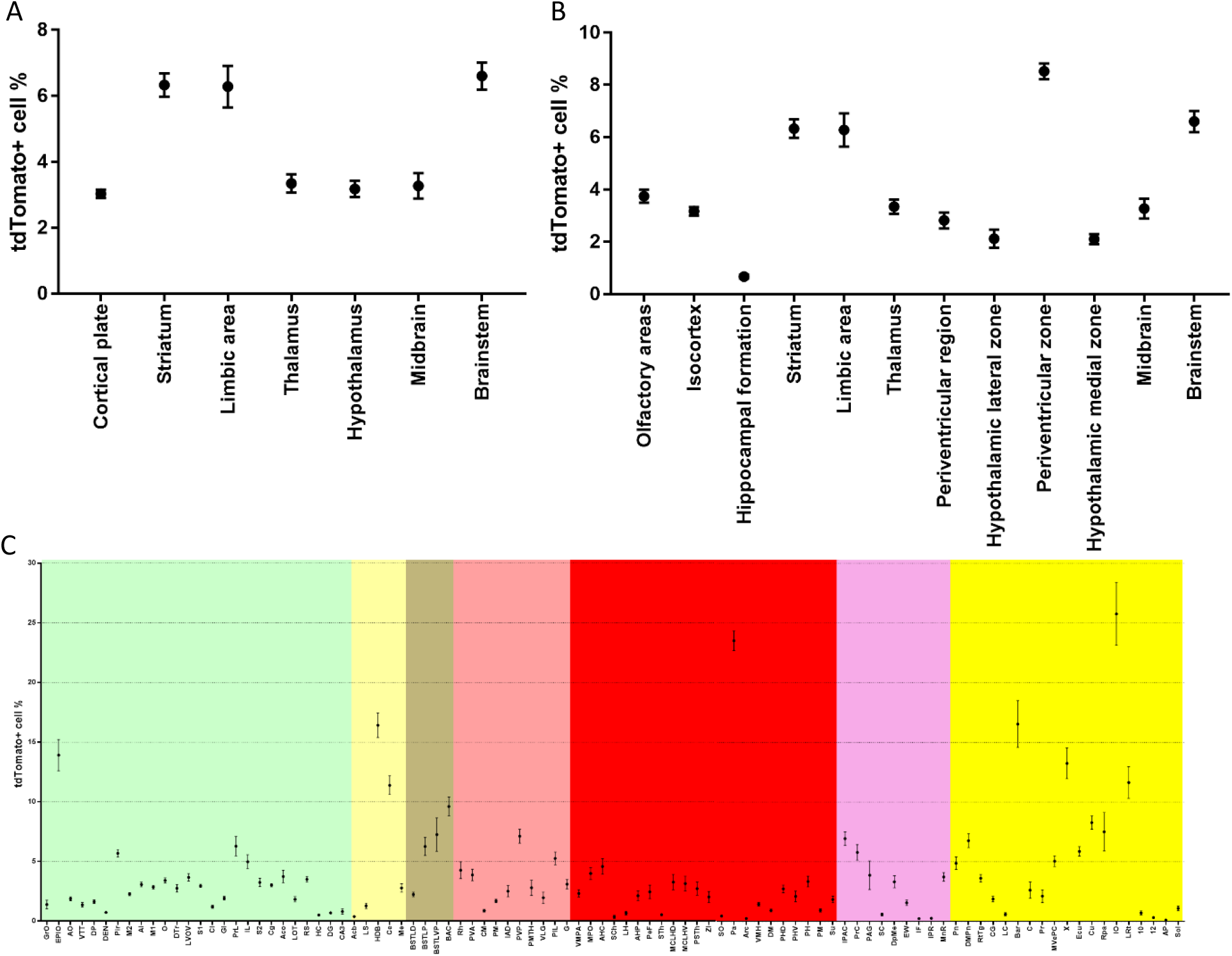
Distribution of the average tdTomato positive cell ratio throughout the CRH-ires-Cre; Ai9 mose brain on the basis of 95 examined regions. Ratio was determined as number of the tdTomato positive cells divided by the total cell amount of the specific region*100. A) Average ratio in cortical plate, striatum, limbic area, thalamus, hypothalamus, midbrain and brainstem partage. B) Average ratio in olfactory area, isocortex, hippocampal formation, striatum, limbic area, thalamus, periventricular region, hypothalamic medial zone, midbrain and brainsem partage. C) Average ratio in each specific reagon.

## 4.3. Neuronal activation

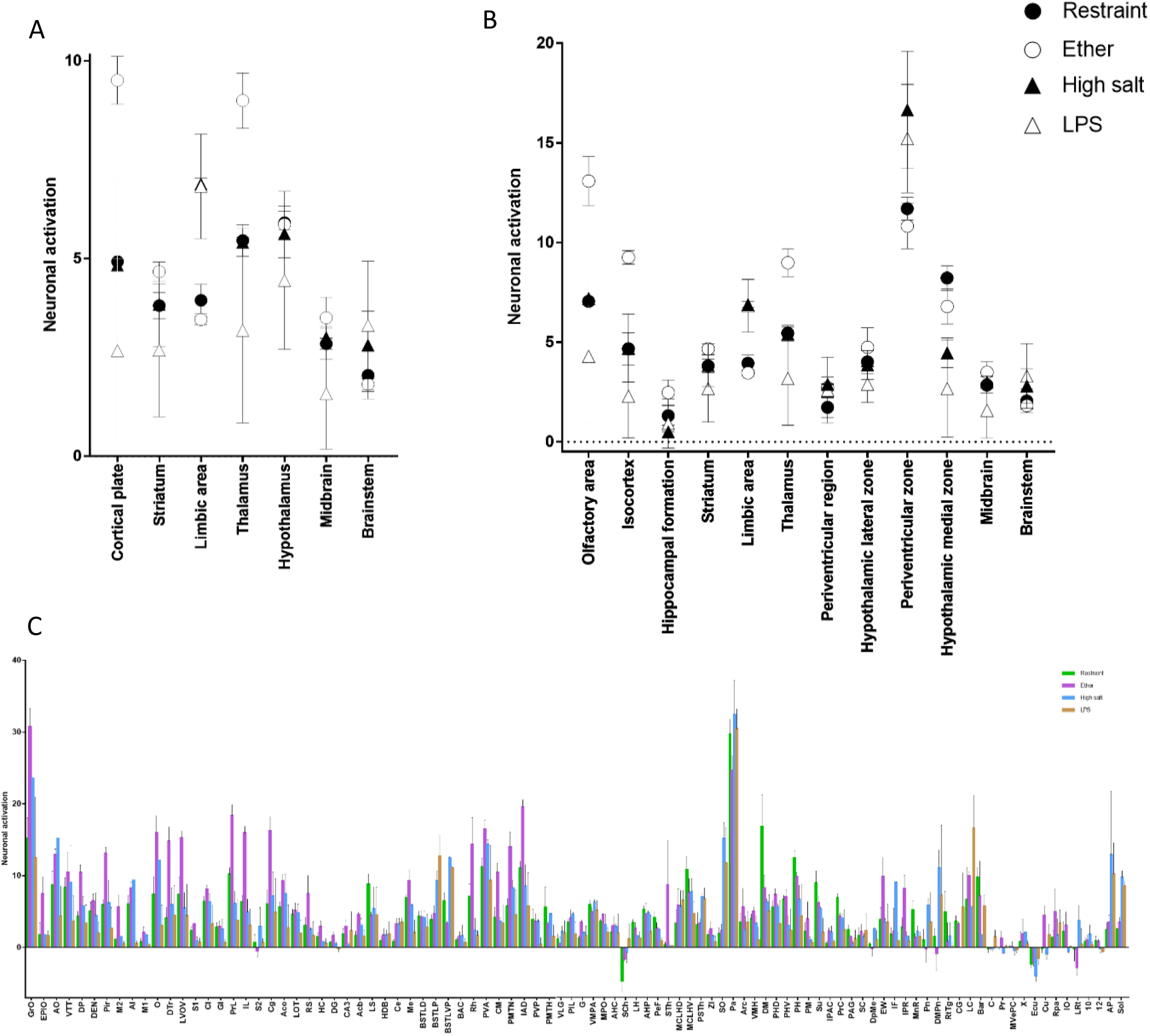
Average neuronal activation alteration of restraint, ether, high salt and LPS exposed animals from the control group. Neuronal activation was determined as treated animal [number of c-Fos positive cells divided by the total cell amount of each region*100] - control animal [number of c-Fos positive cells divided by the total cell amount of each region*100]. A) Average neuronal activation in cortical plate, striatum, limbic area, thalamus, hypothalamus, midbrain and brainstem. B) Average neuronal activation in olfactory area, isocortex, hippocampal formation, striatum, limbic area, thalamus, periventricular region, hypothalamic medial zone, midbrain and brainsem. C) Average neuronal activation in each individually monitored brain region.

## 4.4. Colocalization

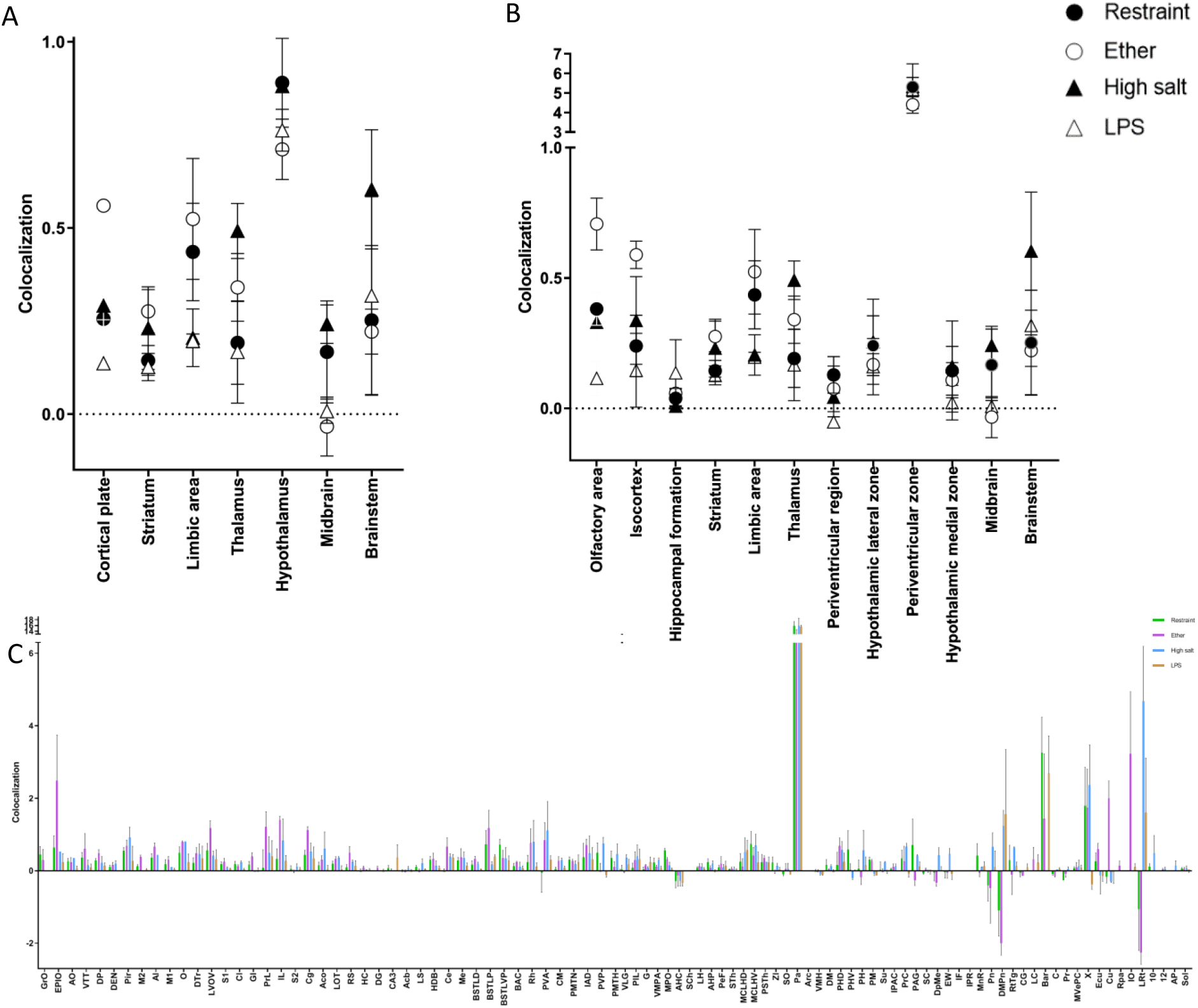
Average colocalization rate change of restraint, ether, high salt and LPS treated mice from the control group. Colocalization rate was determined as treated animal [number of simultaneously tdTomato and c-Fos positive cells divided by the total cell amount of each region*100] - control animal [number of simultaneously tdTomato and c-Fos positive cells divided by the total cell amount of each region*100]. A) Average colocalization in cortical plate, striatum, limbic area, thalamus, hypothalamus, midbrain and brainstem. B) Average colocalization in olfactory area, isocortex, hippocampal formation, striatum, limbic area, thalamus, periventricular region, hypothalamic medial zone, midbrain and brainsem. C) Average colocalization in each monitored brain region.

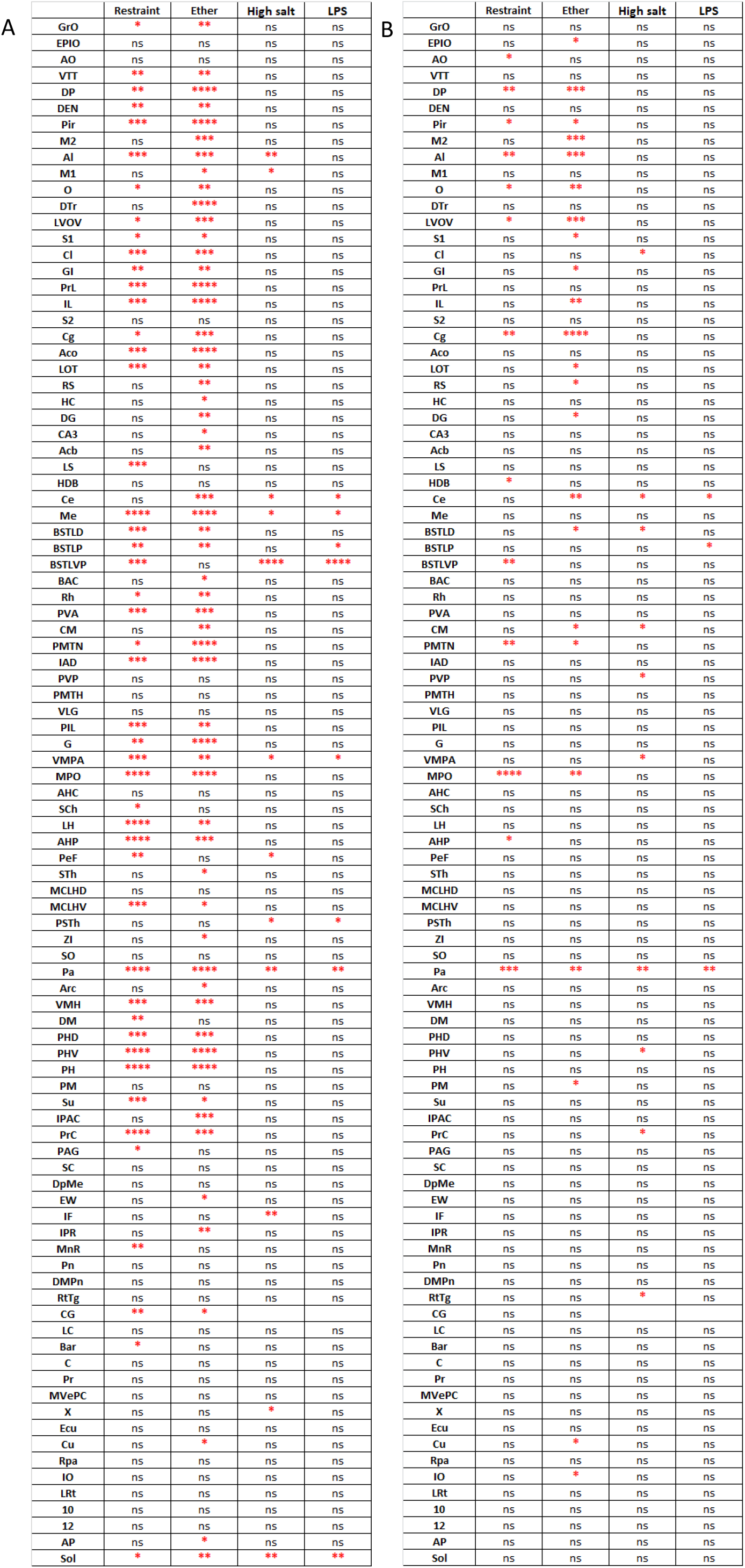
Significance level of discrepancy in A) neuronal activation and B) colocalization rate in case of each treatment group of the 95 examined brain regions. Neuronal activation was determined as number of c-Fos positive cells divided by the total cell amount of each region*100. Colocalization rate was determined as number of simultaneously tdTomato and c-Fos positive cells divided by the total cell amount of each region*100. Datas were analyzed with one-way ANOVA with Tukey’s multiple comparisons test. p*<0.05; p**<0.01; p***<0.001; p****<0.0001; ns=not significant.

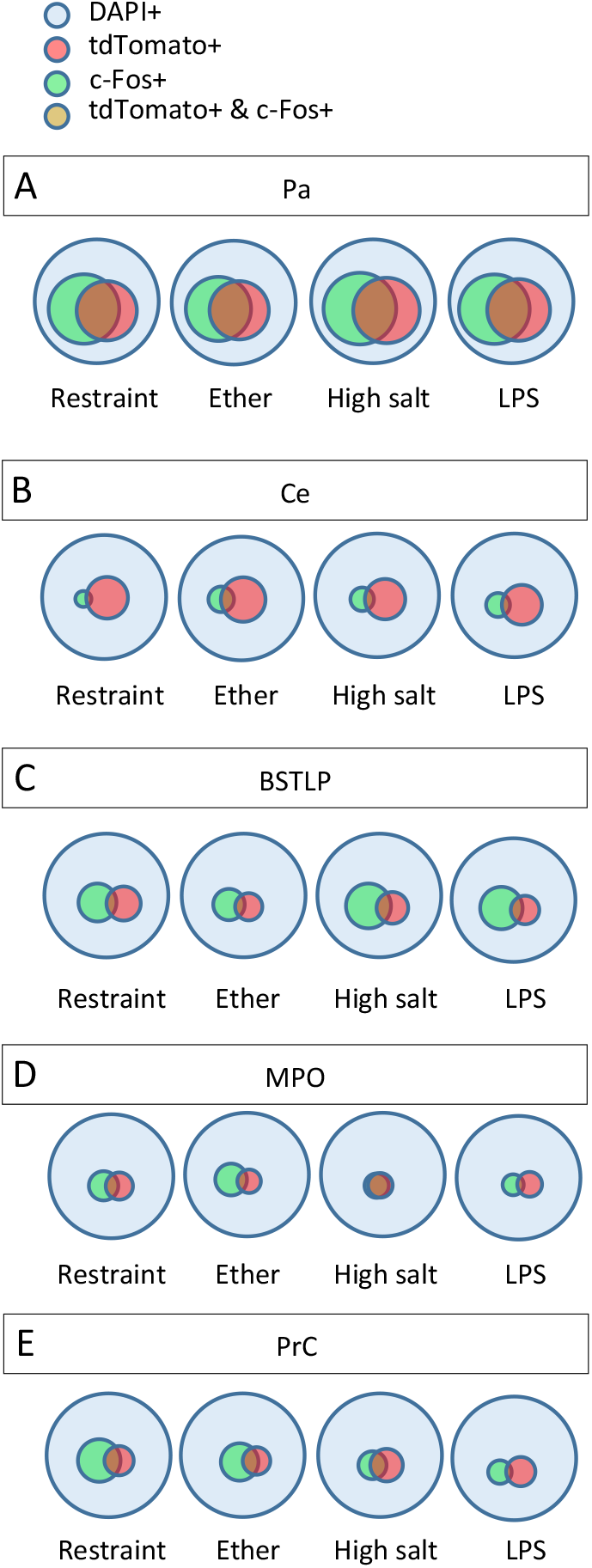
Average proportion of fluorescent signal positive cells in A) Pa; B) Ce; C) BSTLVP; D) MPO and E) PrC. DAPI+ defines total amount of cells; tdTomato+ defines amount of CRH neurons; c-Fos+ defines amount of activated neurons; tdTomato+ & c-Fos+ defines amount of activated CRH neurons.

## 4.5. Stress effect differences

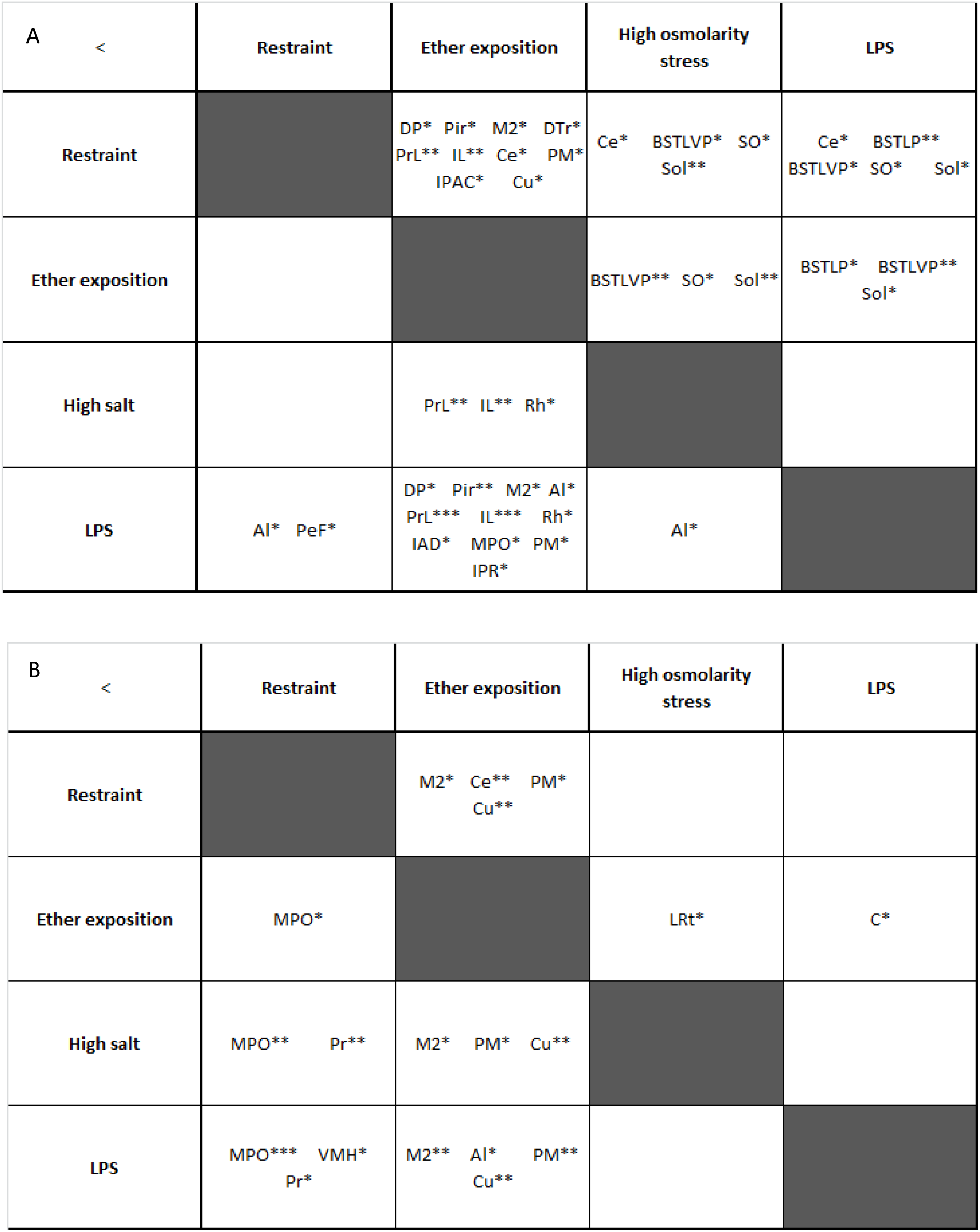
Location and level of significant discrepancy between the stressors in case of A) neuronal activation and B) colocalization growth compared to the control groups. Neuronal activation was determined as treated animal [number of c-Fos positive cells divided by the total cell amount of each region*100] - control animal [number of c-Fos positive cells divided by the total cell amount of each region*100]. Colocalization rate was determined as treated animal [number of simultaneously tdTomato and c-Fos positive cells divided by the total cell amount of each region*100] - control animal [number of simultaneously tdTomato and c-Fos positive cells divided by the total cell amount of each region*100]. Coloumns represent the stressor after which the referred regions gained more neuronal activation or colocalization percentage. Datas were analyzed with one-way ANOVA with Tukey’s multiple comparisons test. p*<0.05; p**<0.01; p***<0.001.

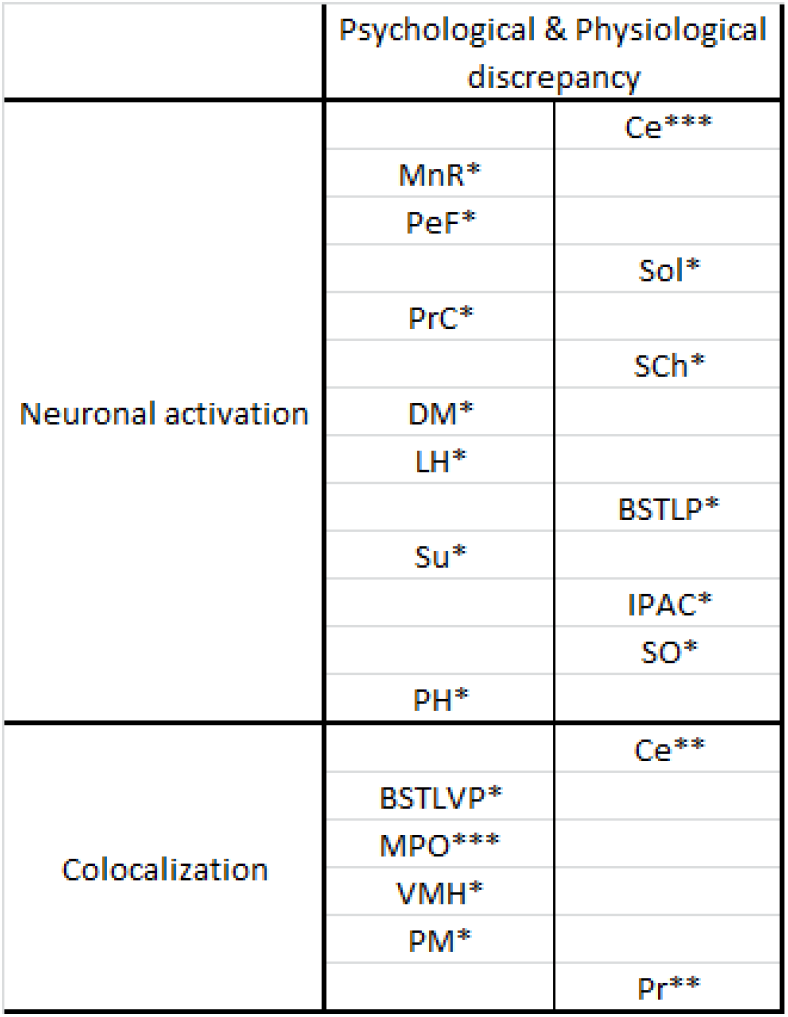
Location and level of significant discrepancy between psychological and physiological stress in case of A) neuronal activation and B) colocalization growth compared to each other. Neuronal activation was determined as treated animal [number of c-Fos positive cells divided by the total cell amount of each region*100] - control animal [number of c-Fos positive cells divided by the total cell amount of each region*100]. Colocalization rate was determined as treated animal [number of simultaneously tdTomato and c-Fos positive cells divided by the total cell amount of each region*100] - control animal [number of simultaneously tdTomato and c-Fos positive cells divided by the total cell amount of each region*100]. Datas were analyzed with unpaired t-test. p*<0.05; p**<0.01; p***<0.001.

## Notes

### Competing Interest Statement

The authors have declared no competing interest.

